# Evolution and manipulation of vector host choice

**DOI:** 10.1101/110577

**Authors:** Sylvain Gandon

**Affiliations:** CEFE UMR 5175, CNRS - Université de Montpellier - Université Paul-Valéry Montpellier - EPHE 1919 route de Mende, 34293 Montpellier, France

**Keywords:** vector borne disease, host choice, parasitic manipulation, mosquito behavior, malaria, virus

## Abstract

The transmission of many animal and plant diseases relies on the behavior of arthropod vectors. In particular, the choice to feed on either infected or uninfected hosts can dramatically affect the epidemiology of vector-borne diseases. I develop an epidemiological model to explore the impact of host choice behavior on the dynamics of these diseases and to examine selection acting on vector behavior, but also on pathogen manipulation of this behavior. This model identifies multiple evolutionary conflicts over the control of this behavior and generates testable predictions under different scenarios. In general, the vector should evolve the ability to avoid infected hosts. However, if the vector behavior is under the control of the pathogen, uninfected vectors should prefer infected hosts while infected vectors should seek uninfected hosts. But some mechanistic constraints on pathogen manipulation ability may alter these predictions. These theoretical results are discussed in the light of observed behavioral patterns obtained on a diverse range of vector-borne diseases. These patterns confirm that several pathogens have evolved conditional behavioral manipulation strategies of their vector species. Other pathogens, however, seem unable to evolve such complex conditional strategies. Contrasting the behavior of infected and uninfected vectors may thus help reveal mechanistic constraints acting on the evolution of the manipulation of vector behavior.

## 1. Introduction

Many animal and plant infectious diseases are transmitted by arthropod vectors. In humans, several deadly vector-borne diseases (e.g. malaria, yellow fever, dengue, West Nile virus) are transmitted by mosquitoes or by other insect species (sandflies, fleas, ticks, tsetse flies). In plants, numerous other vector species (e.g. aphids, leafhoppers, whiteflies) are involved in the transmission of viral and bacterial infections. In spite of the diversity of species involved, the epidemiology of vector-borne diseases can be captured by relatively simple mathematical models describing the pathogen life-cycle across the main host (e.g. a vertebrate, a plant) and the vector (usually an insect). These epidemiological models have clarified the impact of several life-history traits of the vector species for pathogen transmission and pointed out that traits acting on the biting behavior of the vector have a dramatic impact on disease dynamics [1]–[4]. But understanding the evolution of this biting behavior depends on who is controlling this behavior. Indeed, many pathogens are able to manipulate different behavioral traits of their vectors [5]–[7]. Interestingly, the ability of the pathogen to pull the strings of its vector may yield a conflict over the evolution of these traits. For instance, the biting rate maximizing pathogen fitness may be very different from the one maximizing vector fitness [8], [9]. The resolution of this conflict has been studied in several different vector-borne diseases [5]–[7], [10]–[12].

Another important but often overlooked component of transmission involves host choice behavior of the vector. Several experimental studies have demonstrated that some vector species have biased preferences for infected [13]–[16] or uninfected hosts [17]–[19]. Epidemiological models show that vector preference for infected hosts can boost transmission during the early stage of the epidemic [20]–[26]. This suggests that attraction towards infected hosts may result, at least in part, from a manipulation of the vector by the pathogen. Yet, extreme preference for infected (or uninfected) hosts can also limit or even stop pathogen transmission. For instance, if the vectors bite only infected hosts they can never transmit the disease to uninfected hosts. Besides, recent empirical studies in plant pathogens indicate that the host choice behavior may be conditional on the infection status of the vector itself. In particular, uninfected vectors have been found to be attracted towards infected plants but, after being infected, they are attracted towards uninfected plants [27]–[30]. Roosien et al [26] analysed the consequences of these behavioral shifts and demonstrated its dramatic impact for the epidemiology of plant pathogens. Such a conditional modification of vector behavior seems very adaptive for pathogen transmission but this hypothesis remains to be investigated theoretically.

Here I develop a theoretical framework to explore the consequences of vector host choice behavior on the epidemiology and evolution of vector-borne diseases. First, I develop a general epidemiological model to study the impact of the behavior of both infected and uninfected vectors on the persistence of the disease. Second, I use this epidemiological model to study the evolution of vector behavior. Scenarios with or without manipulation are contrasted to discuss the adaptive nature of these modifications of host preference for the vector or for the pathogen. Third, I review experimental studies that have examined host choice behavior in arthropod vectors. In particular, I focus on the handful of studies that have monitored the behavior of both infected and uninfected vectors. The fascinating diversity of vector behaviors in animal and plant vector-borne diseases is discussed in the light of this theoretical model.

## 2. The epidemiological model

Three organisms are interacting in this vector-borne disease model: the host, the vector (usually an insect) and the pathogen (e.g. virus, bacteria, protozoa). The host can either be infected (state *I*) or uninfected (state *S*) and, similarly, the vector can either be infected (state *V_I_*) or uninfected (state *V_S_*). The following set of differential equations governs the dynamics of the densities of these different types of individuals (see table 1 for a summary of the main parameters and the Supplementary Information for details of the derivation of this model):

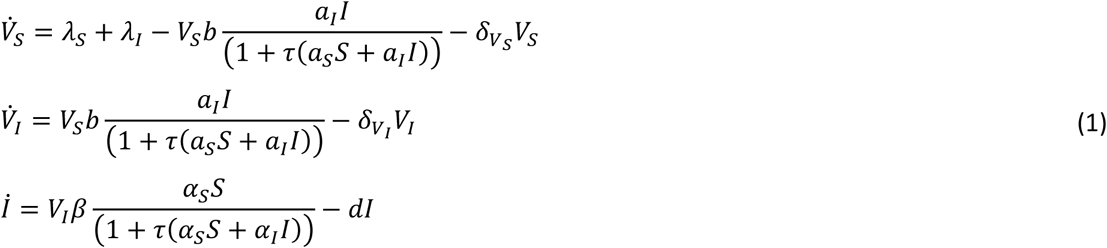

where *λ_S_* = *V_S_f_S_*(1 − *κ N_V_*) and *λ_I_* = *V_I_f_I_*(1 − *κ N_V_*) refer to the density-dependent fecundity of uninfected and infected vectors, respectively. The parameter *κ* measures the intensity of density dependence while *f_S_* and *f_I_* measure the per capita fecundity of uninfected and infected vectors, respectively. The density of the whole population of the vector, *N_V_* = *V_S_* + *V_I_*, is allowed to vary with the dynamics of both uninfected and infected vectors. The first phase of the pathogen life cycle is the infection of the vector after feeding on an infected host. The parameter *b* is the probability that the vector gets infected after biting an infected host. The behavior of the uninfected vectors is governed by the parameters *a_S_* and *a_I_* which refer to the searching efficiency of uninfected and infected hosts, respectively. The parameter *τ* is the handling time of the host by the vector and includes the time taken to bite after landing on the host but also the time taken to digest before an attempt to bite a new host. When the handling time is very small the number of infected bites varies linearly with the number of susceptible hosts. When this handling time is large, it is the frequency of uninfected hosts that governs the epidemiological dynamics [21], [31]. The derivation of this Holling type II response is detailed in the appendix 1. The pathogen is allowed to affect vector survival with specific mortality rates for uninfected and infected vectors (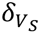 and 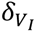, respectively).

The second phase of the pathogen life cycle is the infection of the host by infected vectors. For the sake of simplicity I assume that the total density of hosts, *N* = *S* + *I*, is a constant. This means that whenever a host dies (this occurs at a constant rate *d*) it is immediately replaced by a new susceptible host. The parameter *β* is the probability that the host gets infected after being bitten by an infected vector. The behavior of the infected vectors is governed by the parameters *α_S_* and *α_I_* which refer to the searching efficiency of uninfected an infected hosts, respectively.

**Table 1:** Definitions of the main parameters of the model.

| Main parameters | Definitions |
| --- | --- |
| $N = S + I$ | Host density (susceptible + infected) |
| $N_V = V_S + V_I$ | Vector density (susceptible + infected) |
| $a_S, a_I$ | Searching efficiency of uninfected vectors for uninfected and infected hosts |
| $\alpha_S, \alpha_I$ | Searching efficiency of infected vectors for uninfected and infected hosts |
| $p = \frac{a_I}{a_S + a_I}, \pi = \frac{\alpha_I}{\alpha_S + \alpha_I}$ | Preference towards infected hosts in uninfected and infected vectors |
| $b$ | Probability that an uninfected vector gets infected after biting an infected host |
| $\beta$ | Probability that an uninfected host gets infected after being bitten by an infected vector |
| $\tau$ | Handling time |
| $\lambda_S = V_S f_S (1 - \kappa N_V)$ | Fecundity of uninfected vectors |
| $\lambda_I = V_I f_I (1 - \kappa N_V)$ | Fecundity of infected vectors |
| $f_S, f_I$ | Per capita fecundities of uninfected and infected vectors |
| $F_S, F_I$ | Maximal fecundities of uninfected and infected vectors |
| $\kappa$ | Intensity of density dependence on vector fecundity |
| $d$ | Mortality rate of infected host |
| $\delta_{V_S}, \delta_{V_I}$ | Mortality rates of susceptible and infected vectors |
| $R_0$ | Basic reproduction ratio of the pathogen |
| $R_{Vm}$ | Per generation invasion ratio of a mutant vector |
| $R_{Pm}$ | Per generation invasion ratio of a mutant pathogen |
| $c$ | Cost of searching efficiency on vector fecundity |
| $\phi$ | Quality (for the fecundity of the vector) of the infected host relative to the uninfected host |
| $\sigma_H, \sigma_V$ | Probability of superinfection of an infected host and an infected vector, respectively |

To determine the ability of a pathogen to invade a disease-free environment I derive the pathogen’s basic reproduction ratio *R*_0_ (see appendix 2):

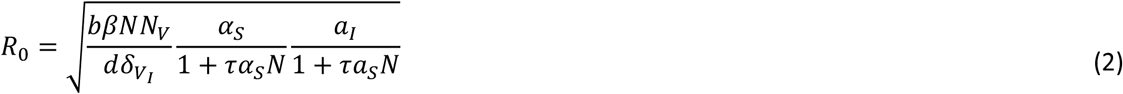

where 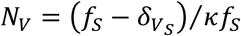 is the equilibrium density of the vector when the pathogen is absent. The pathogen can invade this disease-free equilibrium when *R*_0_ > 1. Higher densities of both hosts and vector are always increasing *R*_0_ but the behavior of both uninfected and infected vectors can also affect the basic reproduction ratio of the pathogen. The preference of uninfected vectors for infected hosts (large *a_I_* and low *a_S_*) and the attraction of infected vectors towards susceptible hosts (large *α_S_*) increase *R*_0_. Note, however, that when *τ* or *N* get very large the basic reproduction ratio depends only on the behavior of uninfected vectors. Under the assumption that the sums of searching efficiencies *a* = *a_S_* + *a_I_* and *α* = *α_S_* + *α_I_* are fixed in uninfected and infected vectors, respectively, one can focus on the effects of the preference between infected and uninfected hosts. More specifically, I introduce the parameters *p* = *a_I_*/(*a_S_* + *a_I_*) and *π* = *αI*/(*α_S_* + *α_I_*) that control the preference towards infected hosts in uninfected and infected vectors, respectively (Table 1). Figure 1A shows that *R*_0_ is maximized when uninfected vectors prefer biting infected hosts and when infected vectors prefer biting uninfected hosts. The figure also illustrates that extreme choice strategies can lead to parasite extinction (i.e. *R*_0_ < 1).

After pathogen invasion the system reaches an endemic equilibrium where the host, the vector and the pathogen can coexist (the notation 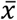 is used to refer to the equilibrium density of the variable *x* at this endemic equilibrium). These equilibrium densities depend on the behavior of the vectors as well as all the other parameters of the model. I failed to find simple analytic expressions for those densities but they can be readily obtained numerically using (1).

Note that the per capita fecundities *f_S_* and *f_I_* were assumed to be fixed quantities in figure 1A. The fecundity of many vector species, however, is likely to be limited by the availability and/or the quality of different types of hosts. Consequently, the fecundity of both infected and uninfected vectors are also going to depend on vector behavior:

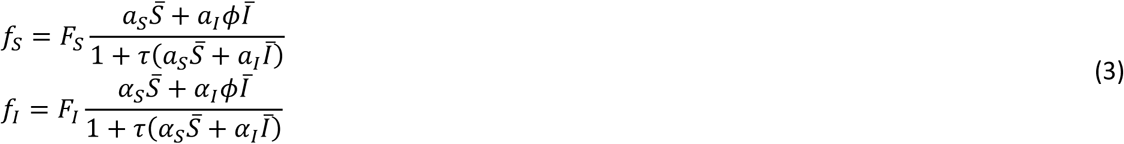

**Figure 1:**
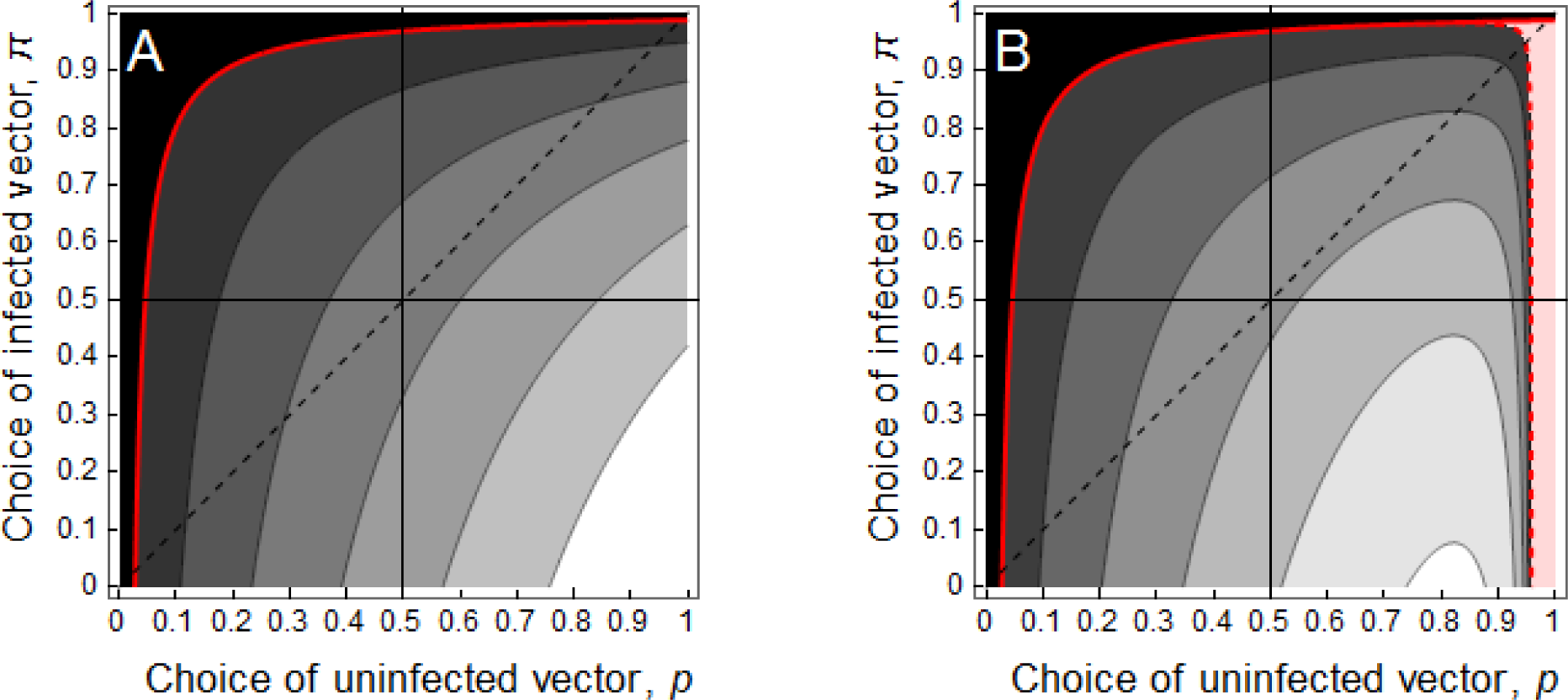
Effect of vector host choice (preference for the infected host) on the basic reproduction ratio Ro of the pathogen given in equation (2). The red line indicates the threshold value where *R*_0_ = 1 and the shades of gray indicate different values of *R*_0_ from 1 to 6 (darkest to lightest). In (A) the population size of the vector, *N_V_*, before the introduction of the pathogen does not depend on host choice behavior because vector fecundity is assumed to be constant *f_S_* = 10. In (B) the population size of the vector, *N_V_*, depends on vector behavior because fecundity is assumed to depend on host preference as indicated in equation (3) with *F_S_* = 10. Note that when uninfected vectors prefer infected hosts, the system exhibits a backward bifurcation at *R*_0_ = 1 (dashed red line) and, depending on the initial conditions of the system, the pathogen may either go extinct or reach an endemic equilibrium when *R*_0_ < 1 (light red region). The full red line and the black area indicate the parameter region where the pathogen is always driven to extinction. Other parameter values: *N* = 500, *κ* = 0.01, *b* = *β* = 1, *d* = 0.05, 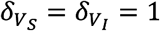, *τ* = 0.1, *a* = *α* = 0.01.

where *F_s_* and *F_I_* are the maximal fecundities of uninfected and infected vectors, respectively. The parameter *ϕ* measures the intrinsic quality of the infected host relative to the uninfected host. For instance, *ϕ* < 1 indicates that infected hosts may provide less nutrients than healthy ones (e.g. in the case of malaria because of anaemia). The influence of vector behavior on vector fecundity can lead to complex epidemiological dynamics. For instance, the dynamical system may exhibit backward bifurcation at *R*_0_ = 1. In other words, depending on the initial condition, the pathogen may either go extinct or reach an endemic equilibrium when *R*_0_ < 1. In particular, this occurs when preference of uninfected vectors towards infected hosts becomes very pronounced (figure 1B). In the following, for the sake of simplicity, I will focus on situations where *R*_0_ > 1.

## 3. Evolution

In the following I study the long-term evolutionary dynamics of the above dynamical system. Using the classical formalism of Adaptive Dynamics I assume mutation rate to be low which allows decoupling evolutionary and epidemiological dynamics [32]–[35]. In other words, I study the evolution of vector behavior (i.e. searching efficiency, host choice preference) through the derivation of the invasion of rare mutants (the subscript *m* refers to the mutant) in a resident system at equilibrium. First I analyze the evolution of vector behavior when this behavior is governed by the vector itself. In a second step I examine a situation where vector behavior is (at least partly) manipulated by the pathogen and evolution takes place in the pathogen population.

### 3.1 Vector evolution

The model can first be used to study the evolution of vector behavior in the absence of the pathogen. In this case all the vectors are uninfected but they can adopt different searching efficiency strategies. Higher searching efficiency allows the vector to exploit more hosts and thus to produce more offspring but, on the other hand, searching for hosts may be costly because more energy is allocated into flying. I analyze the evolution of searching efficiency in appendix 3 and I show that the evolutionary stable searching efficiency decreases with the host population size, *N*, the handling time, *τ*, or the fecundity cost associated with higher allocation to searching efficiency.

When the pathogen is present, the invasion of the mutant vector involves two compartments since the vector can either be infected or not. The invasion of a mutant vector can be analyzed using the per-generation invasion number [36] (appendix 3):

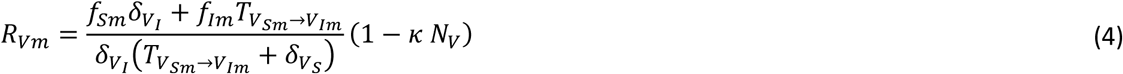

where 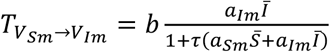.

One could use the above invasion condition to study the evolution of searching efficiency but I want to focus on the preference for uninfected or infected hosts. I will thus assume that the searching efficiencies *a* = *a_S_* + *a_I_* and *α* = *α_S_* + *α_I_* of uninfected and infected vectors are fixed and I will focus only on the evolution of the preference between infected and uninfected hosts. More specifically, I will study the evolution of parameters *p* and *π* that control the preference towards infected hosts in uninfected and infected vectors, respectively (Table 1).

The derivation of evolutionarily stable strategies can be obtained by maximizing *R_Vm_* when the endemic equilibrium (i.e. 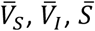 and 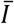) is set by the resident strategy (i.e. *p* and *π*). Factors governing the direction of selection on vector behavior are detailed in appendix 3. In short, the model allows taking into account multiple evolutionary forces: (i) the cost of looking for a rare host, (ii) the cost of feeding on infected hosts, (iii) the potential fitness costs associated with the reduction of the fecundity and/or the survival of infected vectors. In other words, vector evolution is driven by time-limitation (risk of dying before reproducing) and/or egg-limitation (risk of producing a lower number of eggs) as in classical models of life-history evolution of parasitoids [37]. In malaria, for instance, the impact of the infection on vector survival is reduced but it is often associated with a reduced fecundity [38]–[40]. These fitness costs are expected to select vectors that avoid biting infected hosts. But, if the prevalence of infected hosts is very high the opposite may be predicted because the vector cannot afford to lose too much time looking for rare uninfected hosts. For instance figure 2 shows the evolutionary stable strategy of the vector when it is unable to adopt conditional strategies (i.e. *p* = *π*). For a broad range of parameter values the vector prefers to bite uninfected hosts (figure 3A) but when the prevalence in the infection is very high in the host population, the vector may evolve a preference towards infected hosts.

**Figure 2:**
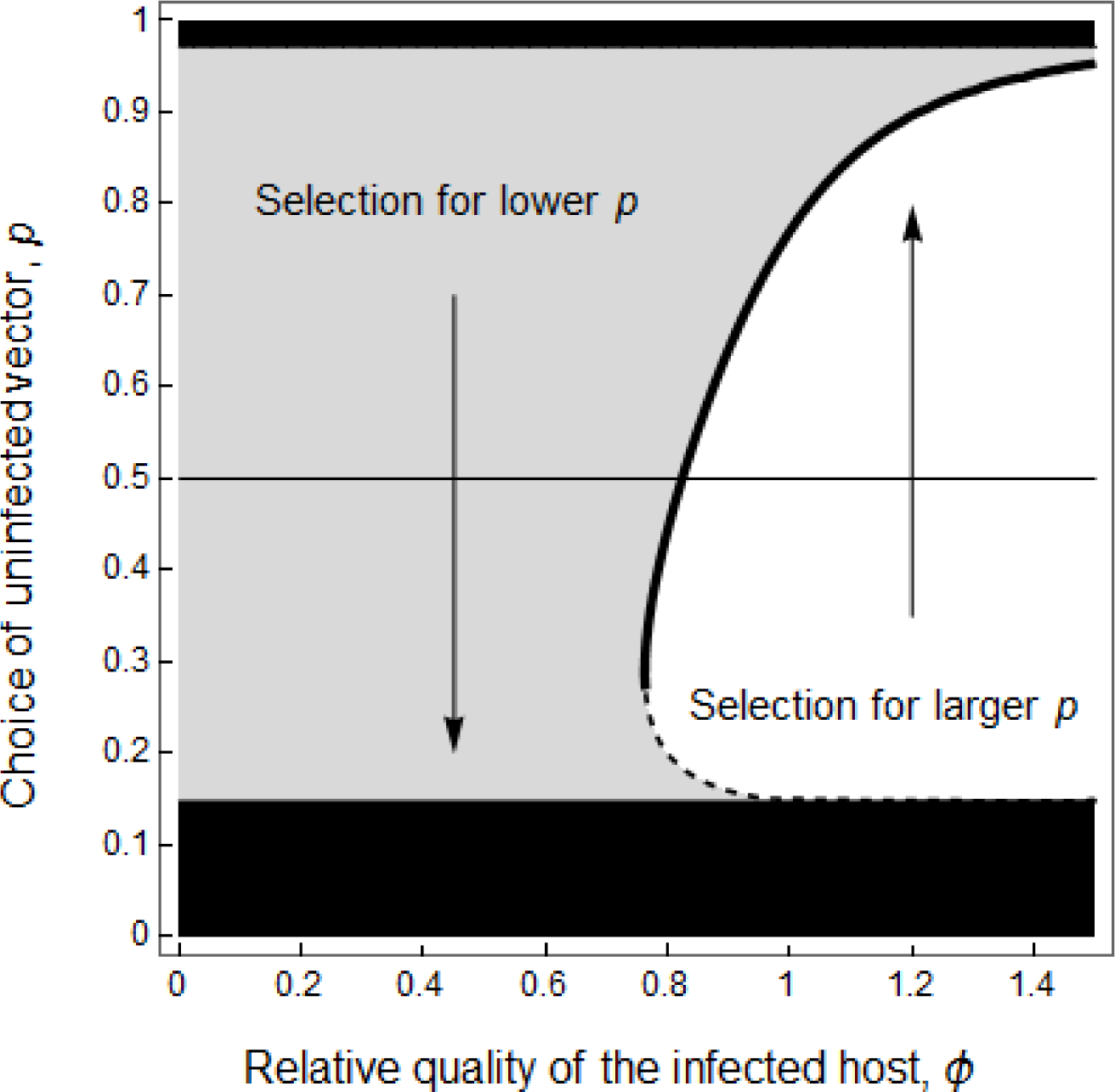
Effect of the relative quality of the infected host, *ϕ*, on the evolution of unconditional host choice (preference for the infected host) of the vector. The pathogen goes extinct when vector preference reaches extreme values (black region). When the relative quality of the infected host is low vectors evolve preference for uninfected hosts (gray region). But when the quality of infected host is relatively high vectors can evolve preference for infected hosts. The ultimate outcome may either be an intermediate preference strategy (bold black line) or extreme avoidance strategy towards infected hosts and, consequently, pathogen extinction. Other parameter values: *N* = 1000, *κ* = 0.001, *b* = *β* = 0.5, *d* = 0.1, 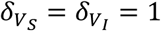, *τ* = 0.5, *a* = α = 0.01, *F_S_* = *F_I_* = 5.

**Figure 3:**
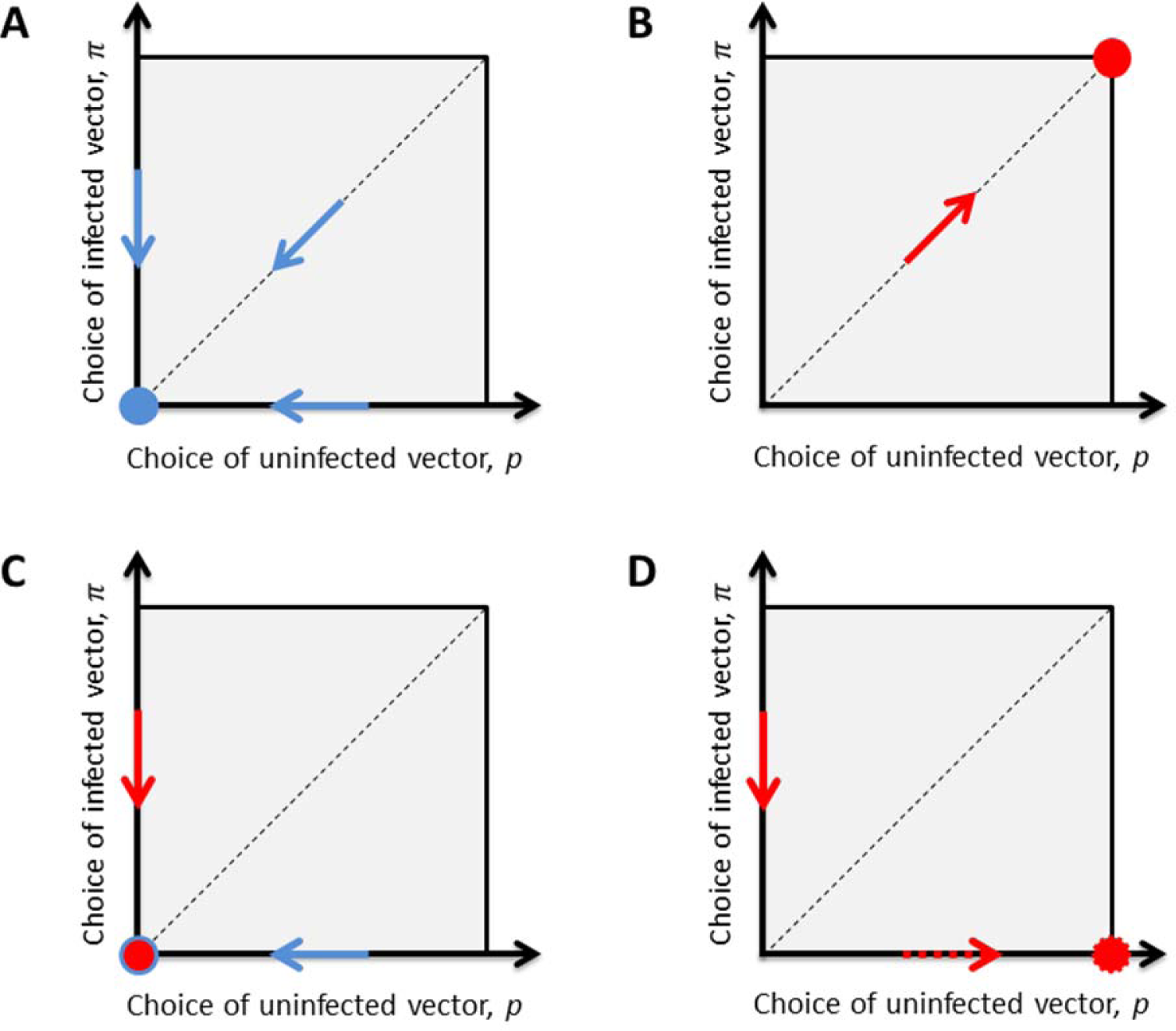
Schematic representation of the evolution of host choice (preference for the infected host) by uninfected and infected vectors under different scenarios. Blue arrows indicate the direction of evolution under vector control and red arrows indicate direction of evolution under pathogen control. In (A) the host choice is only governed by the vector. In (B) the host choice of both infected and uninfected vectors is governed by the pathogen in the infected host (i.e. *p* = *π*). In (C) the host choice of infected vectors is governed by the pathogen while the host choice of uninfected vectors is governed by the vector. In (D) the dashed arrow indicates evolution driven by the pathogen in the infected hosts while the full arrow indicates evolution driven by the pathogen in the infected vector. These four different scenarios yield different ultimate evolutionary outcomes indicated by a large point in each panel. Note that the above panels summarize general evolutionary trends under biologically relevant parameter values but extreme parameter values may yield qualitatively different evolutionary predictions (see main text).

Note, that our analysis yields extreme preference strategies that may ultimately lead the pathogen population to extinction (figure 1). This is because the current model assumes that any preference strategy can evolve. Preference, however, requires an ability to discriminate between different types of hosts. In most biological systems this ability is likely to be imperfect or to carry fitness costs. The above model can be readily modified to account for an intrinsic cost associated with strong preference strategies but this would obscure the qualitative understanding of the evolutionary analysis.

### 3.2 Pathogen evolution

In the above section the vector was allowed to evolve different host preference strategies. But what if these preferences are governed (at least partly) by the pathogen? To answer this question I focus on the dynamics of a mutant pathogen in a resident pathogen population. Using a generalization of classical superinfection models [41], [42] it is assumed that when a vector infected with strain *i* bites a host infected with strain *j* the vector has a probability *σ_V_* to lose the strain *i* and to become infected with strain *j*, while the host has a probability *σ_H_* to lose the strain *j* and become infected with strain *i*. Although this is a very crude approximation of the within-host competition taking place between different pathogens it allows to account for multiple infections in the vector and in the host (e.g. in malaria [43], [44]). The ability of the mutant to outcompete the resident pathogen can be studied using the per-generation invasion number of the mutant (see appendix 4):

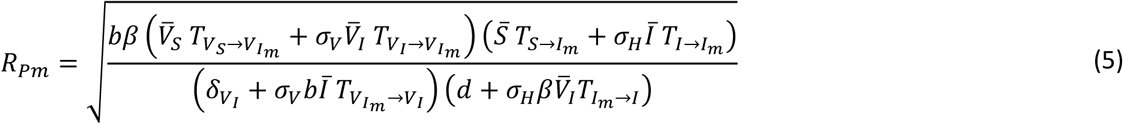

where the notation *T_X→Y_* refers to the transition between the states *X* and *Y*. Importantly, these transitions depend critically on the way the pathogen acts on the behaviour of the vectors. In the following I will consider three different scenarios.

#### Pathogen manipulates vectors from within infected hosts

The pathogen may act on vector behaviour through a manipulation of the attractiveness of the infected host. For example, this manipulation could act through the modification of the volatiles emitted by the infected hosts. In this scenario the behaviour of infected and uninfected vectors are undistinguishable (i.e. *p* = *π*) because both types of vectors are attracted by the volatiles released by infected hosts (figure 3B). In this case the pathogen evolves a manipulation strategy that attracts the vector towards infected hosts. In other words selection on the pathogen is driven by the necessity to attract uninfected vectors even if it also attracts infected vectors. Superinfection in the vector, *σ_V_*, enhances this trend because even already infected vectors can transmit the mutant pathogen currently in the infected host. In contrast, superinfection in the host, *σ_H_*, decreases the magnitude of selection because the mutant currently in the host may be ousted by another strain introduced by infected vectors.

#### Pathogen manipulates only infected vectors

Next, I assume that infected vectors are manipulated by the pathogen from within the infected vector. In the absence of host superinfection the pathogen is always evolving manipulation strategies leading higher vector preference towards uninfected hosts. Superinfection in the vector, *σ_V_*, enhances this trend because the mutant pathogen currently in the vector may be ousted by another pathogen strain if it bites an already infected host. Superinfection in the host, however, may counteract this trend (and may even select preference for infected hosts) because the mutant currently in the vector may outcompete another pathogen strain in an already infected host. Biologically relevant parameter values (i.e. low probability of superinfection, intermediate prevalence) yields preference for uninfected hosts (figure 3C). The behavior of uninfected vectors is driven by selection acting on the vector which yields uninfected vectors to avoid infected hosts. In other words, I recover the prediction obtained when the vector controls its own behavior (compare figure 3A and 3C).

#### Pathogen manipulates independently the preference of infected and uninfected vectors

Finally I consider a situation where manipulation is conditional because it can act both from within infected vectors and from within infected hosts. I only consider the case where the manipulation of infected vectors is fully governed by the pathogen in the vector and the pathogen in the infected host can only affect the behaviour of uninfected vectors. In this case selection favors very different conditional strategies in infected and uninfected vectors. The pathogen manipulates uninfected vectors to bite infected hosts and it manipulates infected vectors to bite uninfected hosts and to avoid infected hosts (figure 3D).

In conclusion the model clearly shows that different assumptions regarding the control of vector behaviour have major consequences on the evolutionary and coevolutionary outcome (figure 3). In particular, I see that if the vector is fully controlling its behaviour it should generally avoid feeding on infected hosts. When this preference is at least partly manipulated by the pathogen three different evolutionary outcomes are possible depending on the mechanisms of the manipulation. These different evolutionary outcomes reveal the existence of conflicts between the vector and the pathogen over the control of vector behaviour. But they also reveal conflicts between the pathogen in the host (who is trying to attract uninfected vectors) and the pathogen in the vector (who is trying to get access to uninfected hosts).

## 4. Experimental studies of host choice behavior

It is particularly interesting to contrast the above theoretical predictions with available information on vector preference in different host-parasite systems. Most of the experimental and empirical work investigating the relative preference for infected or uninfected hosts focused only on host-choice behavior of uninfected vectors. I review this work below before discussing the more limited number of studies that monitored the host-choice behavior of both infected and uninfected vectors (figure 4 and Table S1).

**Figure 4:**
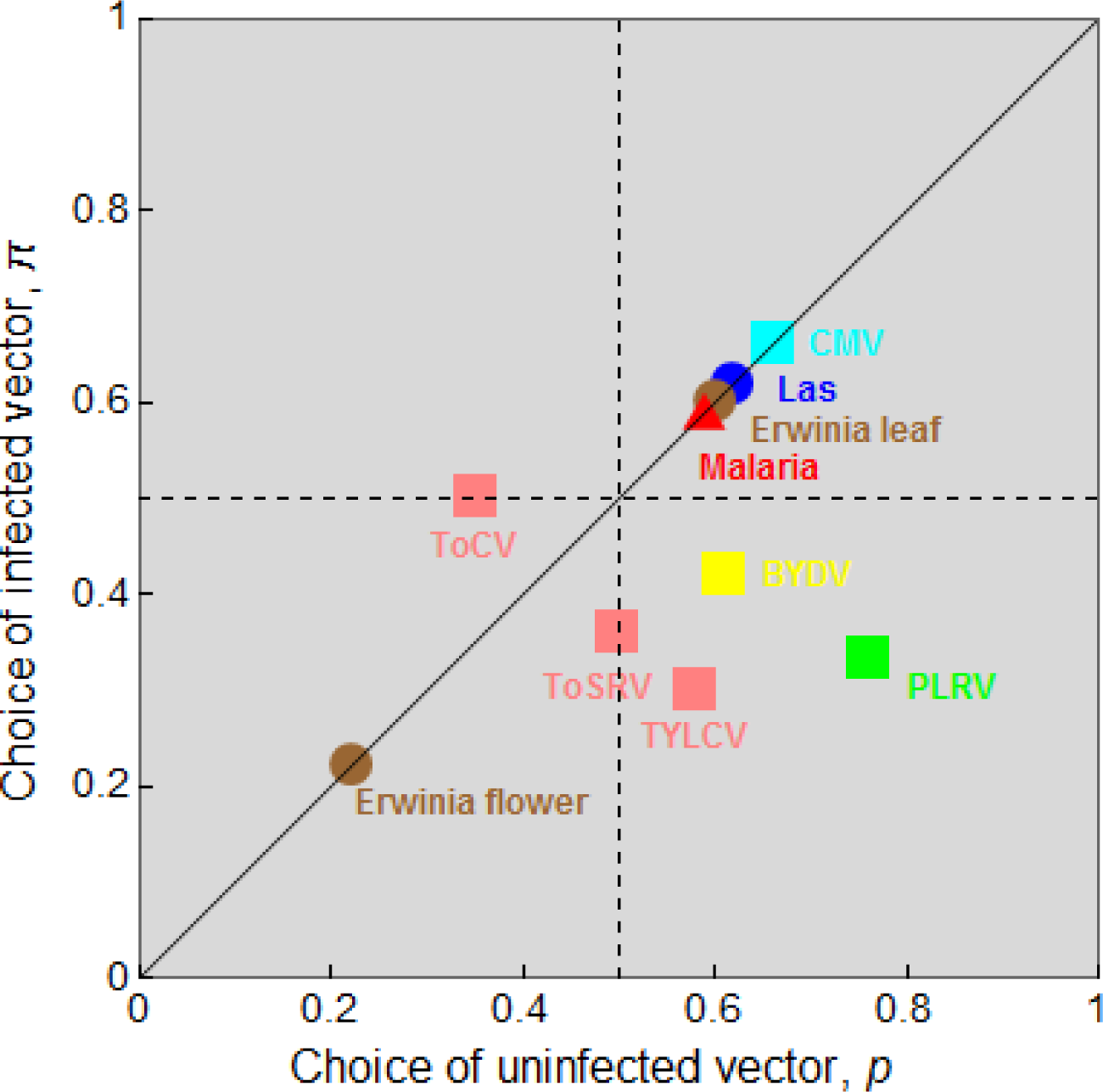
Host choice (preference for the infected host) in uninfected and infected vectors of different pathogens: Tomato yellow leaf curl virus (TYLCV), Tomato severe rugose virus (ToSRV), Tomato chlorosis virus (ToCV), Potato leaf roll virus (PLRV), Cucumber mosaic virus (CMV), *Candidatus liberibacter asiaticus* (Las), *Erwinia tracheiphila* (wilt disease), *Plasmodium relictum* (avian malaria). A detailed presentation of the references used to make this figure is presented in Table S1. Different symbols are used to distinguish between viruses (circle), bacteria (square) and protozoan (triangle).

### 4.1 Behavior of uninfected vectors

First, there is evidence that some vector species have evolved the ability to discriminate and avoid infected individuals. For instance, sharpshooter leafhoppers, a vector of the generalist plant pathogen *Xylella fastidiosa*, are more attracted towards healthy grapevines than symptomatic ones [19]. Similarly, chickens infected with *Plasmodium gallinaceum* have been found to be less attractive to the mosquito vector *Aedes aegypti* [17]. This observed preference for uninfected hosts is likely to be an adaptation of the vector who is trying to avoid low quality hosts (figure 3A). In the Anther-smut disease caused by *Ustilago violacea* the transmission of the spores relies on pollinator visits. Several studies report that insect vectors have the ability to avoid infected flowers [18], [45]. This may result from an adaptation of the vector because infected flowers do not produce any pollen (*ϕ* < 1). But the fact that infected plants are known to bloom earlier and to produce more flowers may attract pollinators early in the season which is likely to result from an adaptation of the pathogen to maximize its transmission to healthy plants later on in the season [45].

Second, several studies found evidence of pathogen manipulation where infected hosts tend to attract uninfected vectors. For instance, hamsters infected with *Leishmania infantum* are more attractive to female sandflies [46]. Humans infected with *Plasmodium falciparum* and mice infected with *Plasmodium chabaudi* are more attractive to their respective mosquito vectors [14], [47]. Phytoplasma are bacterial plant pathogens that are known to convert infected plants into more attractive hosts for their leafhopper vectors [48]. The causative agent of mummy berry disease of blueberry is the fungal pathogen *Monilinia vaccinia-corymbosi* which induces the production of pseudoflowers and mimicry of floral volatiles that attract insect vectors towards infected plats [49].

### 4.2 Behaviour of infected and uninfected vectors

Most of the work on conditional preference in vectors has been carried out in plant pathosystems. One of the first studies testing for such conditional preference has been done with Barley yellow dwarf virus (BYDV). Although the noninfected aphid *Rhopalosipum padi* prefers to feed on infected wheat plants, the acquisition of the virus dramatically alters the behavior of the infected vector that prefers noninfected plants [27]. A similar reversal of feeding preference has been found in aphids infected by Potato leafroll virus (PLRV) [50]. This preference is mediated by virus-induced changes of potato plants that emit volatile blends enriched in monoterpenes, aldehydes and sesquiterpenes [50]. The Tomato yellow leaf curl virus (TYLCV) can also alter the host preference of its whitefly vector in the same way but, interestingly, this switch is modulated by the genotype of the vector, the genotype of the host and the timing of the infection [51], [52]. In particular the conditional preference of the whitefly was prominent only 6 weeks after infection on susceptible genotypes of plants. The Tomato severe rugose virus (ToSRV) is another virus infecting tomatoes where viruliferous (i.e. infectious) whiteflies are attracted towards volatiles emitted by uninfected plants, while non-viruliferous whiteflies do not show any preference between volatiles emitted by infected or uninfected plants [53]. These four different plant viruses match the above theoretical prediction when pathogen manipulates independently the preference of infected and uninfected vectors (figure 3D).

Bacterial pathogens have also been found to affect the behavior of insect vectors. For instance, *Candidatus liberibacter asiaticus* (Las) has been found to enhance attraction to both uninfected and infected psyllid vectors [54]. This differential preference appears to be mediated by pathogen-induced emission of methyl salicylate in infected plants. The fact that infected vectors also prefer infected plants is likely to result from the emission of a general signal that attracts all the aphids (uninfected or not infected vectors) as discussed in our theoretical model (figure 3B). Note, that the pathogen has evolved other ways to encounter uninfected plants. First, aphids landing on infected plants are rapidly driven away from these poor quality hosts (lower palatability). Second, infected aphids have been found to increase their propensity to disperse (i.e. higher *α* in our model) which may also allow the pathogen to settle in new uninfected plant populations. The bacterial pathogen *Erwinia tracheiphilia* [55] was also found to alter the foliar and floral volatile emission of its wild gourd host in ways that attract the beetle vectors towards infected leaves and uninfected flowers. The beetles may thus acquire *E. tracheiphilia* from infected leaves and transmit the pathogen to a new plant because cucumber beetle via attraction of their uninfected flowers. This differential effect on leaves and flowers may be yet another way to promote pathogen transmission between infected and healthy plants even though infection of the beetle does not seem to affect its preference.

All the above examples refer to plant pathogens that reside for extended periods in their vectors. These persistently transmitted pathogens (PTP) have therefore more opportunities to act on the preference of their insect vectors. In contrast, some plant pathogens are non-persistent in their vectors (NPTP) and are expected to have lower abilities to act on vector behavior [29]. For instance the *Cucumber mosaic virus* (CMV) bind to specific regions of the mouthparts of the vector and are acquired and inoculated during brief tastes of outer plant cells. CMV has been shown to increase the volatile emissions in infected plants and to attract aphid vectors [29]. But CMV also alters nutrient cues of infected plants and this reduction of palatability encourages aphids to seek new and possibly uninfected plants. This pathogen manipulation is likely to enhance pathogen transmission but the conditional change of vector preference is driven by the poor quality of the infected host which is perceived by the vector after landing and not by a direct effect of the pathogen in the individual vector. The Tomato chlorosis virus (ToCV) is semi-persistent but does not circulate in its whitefly vector and seems to induce maladaptive modifications of its vector. Non-viruliferous whiteflies prefer the volatiles emitted by uninfected plants but viruliferous vectors do not exhibit any preference [53].

As far as I know a very limited number of studies have been done on the conditional behavior of infected and uninfected vectors of pathogens of animals. Unlike earlier results obtained with *P. gallinaceum* [17], uninfected *Culex pipiens* mosquitoes are attracted by passerine birds infected with *Plasmodium relictum* [15]. In a subsequent study Cornet et al. [56] showed that both infected and uninfected *C. pipiens* mosquitoes are attracted towards infected birds. This result is in line with the above theoretical predictions when the pathogen manipulates vectors from within the infected hosts (figure 3B). This suggests that the manipulation of host-choice behavior by *P. relictum* acts on the quantity and/or the quality of volatiles emitted by infected birds [47] and that both infected and uninfected mosquitoes are attracted by the scent of this infection. Further studies are required to confirm this prediction and to better characterize the underlying mechanism acting on mosquito behaviour in other malaria parasites including human malaria [57].

## 5. Discussion

The epidemiology of vector borne disease is very sensitive to the host-choice behavior of the arthropod vector. I developed a general model of vector borne transmission taking into account key features of the ecology of a broad range of different pathosystems. Interestingly, this model allows to escape the classical dichotomy between density and frequency dependent models [31] and may help provide a more realistic description of the transmission process of vector borne diseases. This model shows that extreme choice strategies can have dramatic consequences on the epidemiology of the disease and can even lead to pathogen eradication. However, when the uninfected vectors are more attracted towards infected hosts the dynamical system may exhibit backward bifurcation at *R*_0_ = 1. In other words, a stable endemic equilibrium may exist even if *R*_0_ < 1. This result implies that vector choice may prevent the eradication of pathogens even if human interventions managed to reduce *R*_0_ below its critical level. Similar bistability has been observed in models of malaria transmission [58], [59] but here I show that the behavior of uninfected mosquitoes is a key driver of this dynamic. Further work is required to better identify conditions promoting this epidemiological bistability.

The evolutionary analysis of this model reveals complex conflicts between the vector and the pathogen over host-choice behaviour. Under some scenarios, the evolutionary interests of the vector and the pathogen are aligned which leads to a unique evolutionary outcome. In particular, when the pathogen is only able to manipulate the behaviour of infected vectors, both the vector and the pathogen are generally evolving a preference towards uninfected hosts (figure 3A and 3C). In this situation it is impossible to determine who is controlling the evolution of vector behaviour from observed preference patterns.

But pathogen evolution and vector evolution can yield qualitatively very different strategies under other scenarios. This conflict emerges as soon as the parasite in the infected hosts is able to govern host-choice behaviour of the vector. In this case, pathogen selection favours manipulation strategies leading uninfected vectors to prefer infected hosts (figure 3B). Indeed, numerous empirical studies show that pathogens can modify the scent of infected hosts to attract vectors [29]. This manipulation often involves the elevation or exaggeration of existing cues used by vectors to locate hosts. As pointed out by Mauck et al. [60] the evolution of such a “supernormal stimulus” does not involve major qualitative differences between infected and uninfected hosts and it is thus very difficult for the vector to evolve avoidance strategies even if infected vectors suffer from major fitness costs.

Finally, when the pathogen is able to adopt a different strategy in the host or in the vector, conditional preference strategies can evolve. Indeed, the transmission of the pathogen is maximised when uninfected vectors are attracted towards infected hosts and when infected vectors are attracted towards uninfected hosts (figure 3D). Interestingly, only plant viruses with a persistent and circulative mode of vector transmission have been shown to evolve such conditional preference strategies (figure 4). This suggests that only pathogens that evolved a persistent and intimate relationship with their vector are able to induce conditional preference strategies. Note, however, that in spite of persistent infection in its mosquito vector, *Plasmodium* does not induce conditional preference strategies [56]. Further studies exploring the host-choice preference of both infected and uninfected vectors of other pathogens are required to confirm that only virus have the ability to evolve such complex conditional manipulation strategies. In addition, it would be interesting to see if some pathogens are able to evolve other forms of conditional manipulation of host preference varying with the age of the infection in the vector. Indeed, one may expect different manipulation strategies in infected but not yet infectious vectors.

An interesting extension of this work would be to analyse situations where multiple pathogens share the same host and/or the same vector. It is easy to imagine how these complex epidemiological scenarios could yield new evolutionary conflicts over the manipulation of vector behaviours [61]. In addition, many pathogens can infect multiple host species and the vector preference for different host species can also have massive epidemiological consequences [62]–[64]. Several recent studies indicate that preference for different host species is heritable and could thus evolve as a response to a change of the environment [65], [66]. The above theoretical framework could be used to understand and predict the evolution and the manipulation of this other important behavioural trait.

The predictive power of these evolutionary models hinges upon our knowledge of the constraints acting on these behavioural traits. To understand these constraints it is important to study the mechanisms underlying vector preference. Experimental studies on vector preference indicate that host-choice can be mediated by multiple cues like odour, colour and taste [29], [67], [68]. Some pathogens have been shown to modify vector behaviour through the modification of these cues [29], [13], [69]. But in most cases the underlying mechanisms driving these modifications of vector behaviour remain elusive. A better understanding of these underlying mechanisms could also lead to the development of novel public-health strategies to control vector-borne diseases [70]–[73]. The above theoretical analysis provides a framework to understand the evolution and the manipulation of key behavioural traits of vectors (e.g. host choice, biting rate) as well as a guide to structure the exploration of the mechanistic constraints acting on this evolution in a broad range of vector-borne diseases.

## Acknowledgements

I am very grateful to Ana Rivero, Nicole Mideo and Samuel Alizon for comments on the manuscript. This research was funded by European Research Council starting grant 243054 “EVOLEPID”.

